# EZ-DMS – A Simple and Accessible Protocol and Software Package for Deep Mutational Scanning of Virus Proteins

**DOI:** 10.64898/2026.01.02.697422

**Authors:** Aaron N Gillman, Madeline M Broghammer, Samuel A McCarthy-Potter, Rohith Rao Vujjini, Cassian M Birler, Alexander B Kleinpeter, Hillel Haim

**Affiliations:** Department of Microbiology and Immunology, The University of Iowa, Iowa City, IA

**Keywords:** Deep Mutational Scanning, Saturation Mutagenesis, Virus mutational landscape, Fitness Profiling, HIV-1, Envelope Glycoproteins, Antiviral Therapeutics, Temsavir

## Abstract

Mutations that occur during viral genome replication can result in new phenotypes, such as infection of new cell types or resistance to therapeutics. To determine the phenotypic potential of viruses, deep mutational scanning (DMS) can be used. This approach measures the effects of all amino acid changes at sites of interest on protein function, virus infectivity, or resistance to therapeutics. A virus library that contains all possible variants is produced and used to infect a culture of cells. The frequency of each amino acid at the DMS site in the infected culture relative to the input stock captures the relative fitness for each form under the selective pressure applied. Current DMS protocols are complex and costly due to reliance on deep sequencing for genomic characterization. They also require expertise in bioinformatics to process and analyze the data, thus limiting the number of labs that can apply the technology. Here, we describe a simple and efficient protocol to perform DMS on single sites and on two-site combinations in the Env protein of HIV-1. Dual-site DMS allows probing for epistatic effects. The protocol greatly reduces recombination of the proviral genome in transformed bacteria, thus efficiency of the method. Sequencing is performed using Oxford Nanopore technology, and the data are uploaded to a graphical user interface-based platform that calculates the amino acid preferences, which describe the relative fitness of all amino acids under the experimental conditions tested (e.g., under selection pressure of a drug). The protocol does not require expertise in bioinformatics and can be completed in approximately 16 days, with 3-4 days of laboratory activity and 12-13 days of incubation, with considerably lower costs than existing systems.

## INTRODUCTION

Viruses have a tremendous capacity to diversify within the infected host and, consequently, within the population (1–3). The error rate of most viral polymerases ranges between 10^-5^ and 10^-6^ errors per nucleotide per virus replication cycle for RNA viruses and between 10^-6^ and 10^-8^ for DNA viruses (4, 5), resulting in up to one change in each new genome generated. Nucleotide substitutions, deletions and insertions are introduced in coding and non-coding genes, which can reduce virus infectivity. However, they can also impart the virus with new or enhanced features that allow it to persist in the host, including evasion of immune responses (6, 7), infection of new cell types (8, 9), and resistance to antiviral therapeutics (10). To determine the effects of specific changes on the virus phenotype, the mutations can be introduced individually in the gene of interest by site-directed mutagenesis. Properties of the mutant relative to the wild-type form shed light on structure-function relationships, pathogenic potential, and the breadth of therapeutics. While testing of individual mutations is informative, usually more than one is considered, rendering the approach laborious and providing a restricted view of the effects of specific changes.

For a broader understanding of the phenotypic potential of the virus, the effects of all amino acid changes at sites of interest can be tested. The approach, designated saturation mutagenesis or Deep Mutational Scanning (DMS), involves introduction of degenerate codons at the sites. A heterogenous population of viruses (designated a library) is produced that contains all possible non-synonymous substitutions at the positions of interest. The virus library is used to infect a culture of cells. The variants that complete the stage of infection analyzed (e.g., virus genome replication) are detected by deep sequencing of the sample. The output is the amino acid preference profile, calculated as the frequency of each amino acid variant at the DMS site in the infected cells corrected for its frequency in the input virus population. The relative abundance of each variant captures its relative fitness to complete the step of infection analyzed. When conducted in the presence of a therapeutic, this frequency indicates the relative resistance to the agent, and the overall preference profile describes the escape potential of the virus. Multiplexing of libraries, each containing a degenerate codon at a different site, within the same experiment allows testing of up to hundreds of positions in the same protein simultaneously (11, 12).

Saturation mutagenesis has been used for more than two decades to probe the phenotypic potential of proteins from non-viral systems. One of the first applications was to optimize the binding efficiency of antibodies to their targets by introducing degenerate codons in their variable regions, followed by display on the surface of phages (13, 14), yeast (15), or mammalian cells (16). Saturation mutagenesis has also been used to investigate the function of proteins, their interactions with other proteins, and resistance to therapeutics (17) in both bacterial systems (18–20) and in yeast (21–23). Given the ability of viruses to diversify and adapt rapidly to their environment, DMS has also been extensively employed to map the fitness landscape of their proteins. Viruses from diverse families have been profiled using DMS, including influenza virus (24–29), hepatitis C virus (30), Lassa virus (31) and SARS-CoV-2 (11). In addition, human immunodeficiency virus type 1 (HIV-1) has been studied, with specific emphasis on the viral envelope glycoproteins (Envs), the most variable protein of this pathogen. DMS has been applied to profile the fitness landscapes of Envs from diverse strains (12), their resistance profiles to antibody therapeutics (32, 33), and to determine the specificity of antibodies in blood samples from HIV-infected individuals (34). Here, we describe the application of DMS to analyze the Env protein of HIV-1. Nevertheless, as detailed below, the molecular approach and computational tools provided can be directly applied to other HIV-1 proteins as well as other viruses. The specific requirements for such systems are indicated in the Discussion Section.

To generate libraries of HIV-1 particles containing Env variants, two DMS strategies have been previously applied. The classical approach involves the use of replication-competent HIV-1, whereby degenerate codons are introduced in a plasmid that encodes the full-length proviral genome. The advantages of such an approach are: **(A1)** Properties of replicative viruses differ from those of replication-defective viruses, including the number of Env trimers incorporated in viral particles, their processing efficiency, and fusion dynamics (35–37). **(A2)** Replicative viruses can be used during extended infections, or over multiple rounds of infection, to determine the competitive advantage of the variants under sustained selective pressures. The disadvantages of replicative virus systems are: **(D1)** Full-length HIV-1 genomes are unstable in proviral form – replication of such plasmids in bacteria is associated with frequent recombination events in the long terminal repeat (LTR) regions, leading to excision of much of the genome from the construct. Most bacterial cells that are used for this purpose, such as the Stbl *E. coli* cell line (38, 39), only modestly reduce recombination at the LTRs. **(D2)** The use of replication-competent virus requires infections to be performed under biosafety level (BSL) 2+ conditions.

As an alternative to replicative HIV-1, DMS can also be performed using single-round lentiviruses (40). In this system, a library of plasmids that encode for the protein is used to generate a library of cells that express the viral protein library. Viruses are then rescued from the cells by co-transfection with plasmids that express the HIV-1 proteins required for virion production. The replication-defective virus stock is harvested and used to transduce cells, which are then analyzed as in the first approach, by deep sequencing to identify the transducing forms. The main advantages of this system are: **(A1)** Experiments can be conducted under BSL-2 conditions. **(A2)** The cell library, if stable, can be expanded and maintained for multiple experiments. The main disadvantages of this approach are: **(D1)** Single-round pseudoviruses do not capture the stoichiometric relationship between the different constituents of a virus and may thus not reflect well the biology of viruses produced in the host, or their sensitivity profiles to inhibitors (35). **(D2)** The method is limited to the use of virus-producing cells that can be efficiently transfected to allow rescue of the virus. As such, it is based on immortalized cell lines, most commonly human embryonic kidney (HEK) 293T cells. Given the effects of producer cell type on post-translational processing of viral proteins and their resistance profiles to antibodies (41), this method does not accurately represent properties of viruses produced in primary (or immortalized) T-cells. **(D3)** Virus stocks cannot be further cultured for competition assays or analysis of second-order mutations that may emerge.

The two DMS strategies described above share several shortcomings. First, the complexity of the multi-step molecular protocols and of the computational analyses have effectively limited their application to a small number of labs across the world. Indeed, while the investigators have rendered the code publicly accessible (32), the processing of deep sequencing data and code implementation require expertise in bioinformatics that is not available in many molecular biology labs. Second, the sequencing methods usually applied for these protocols are associated with labor-intensive steps and high costs. For short-read sequencing approaches (e.g., Illumina), sample processing involves fragmentation or PCR amplification of short genomic regions followed by index and adapter ligation steps, usually with multiplexing of dozens of indexes for cost efficiency. Third, these approaches require specialized skill sets and costly consumables (*e.g.* library preparation kits and flow cells) that can be prohibitive for smaller labs or narrowly focused studies. Lastly, HIV-1 genomes are unstable in proviral form, and replication of plasmids in bacteria is associated with frequent recombination events in the long terminal repeat (LTR) regions, leading to excision of much of the genome from the construct. Most bacterial cells that are used for this purpose, such as the Stbl *E. coli* cell line (38, 39), only modestly reduce recombination at the LTRs and exhibit poor transformation efficiency due to the large size of the vectors (>14 kb). To address these shortcomings, we developed the EZ-DMS protocol and associated software suite.

Given the broad utility of DMS to understand the evolutionary space of diverse viruses, we sought to increase the accessibility of this method to the broader scientific community. To this end, we developed a new step-by-step protocol with associated software to perform DMS on a single site or dual sites using replication-competent HIV-1. In the development of this protocol, our first goal was to simplify the cloning of the fragment that contains the degenerate codons into the provirus-expressing vector. To this end, instead of using a restriction enzyme-based approach, which requires unique sites in the vector backbone, we used the recombination-based In-Fusion cloning method (Takara Bio), as we described (1). The approach is simple, efficient, and allows the same vector backbone, linearized once, to be used repeatedly for studies across the same gene or region.

Our second goal was to reduce the recombination of the plasmids at the LTRs while maintaining adequate transformation efficiency, plasmid yields, and plasmid diversity. To this end, we used Zymo Mix&Go! DH5α cells produced with the Zymo Mix & Go! *E. coli* Transformation Kit. These cells are distinct from common DH5α cells in that they contain the deoR mutation, which promotes uptake and stability of large plasmids. The transformed bacteria are plated without outgrowth, preventing plasmid propagation during recovery and ensuring that each colony is an independent transformation event. By eliminating the typical recovery and outgrowth phase following transformation, we also nearly eliminated recombination events at the LTRs. Furthermore, by utilizing the Mix & Go! *E.* coli Transformation Kit, we produced Mix & Go! Competent DH5α™ cells at a fraction of the cost of alternative commercial cloning strains.

Our third goal was to apply an inexpensive and labor-non-intensive sequencing approach. To this end, we used the Oxford Nanopore (ONP) long-read sequencing platform. An advantage of ONP sequencing is that it does not require ligation of adapters as in short-read sequencing approaches and can provide sequence data for fragments up to 25,000 nucleotides in length. Furthermore, the high accuracy of current ONP data processing algorithms (42) and low cost rendered this approach favorable.

Finally, our fourth goal was to construct a simple executable graphical user interface (GUI)-based system for processing the sequence data, which outputs the corrected amino acid preference for each site and accurately partitions data from multiplexed DMS experiments of different positions.

## MATERIALS AND METHODS

The protocol described in this peer-reviewed article is published on protocols.io, updated December 31, 2025, https://doi.org/10.17504/protocols.io.n2bvje19wgk5/v1 and is included for printing as Supporting Information S1 File with this article.

### (1) Plasmid Library Production

As shown in **Figure 1**, production of the plasmid library that contains all amino acid variants at a selected position is divided into the following steps:

**Figure 1:**
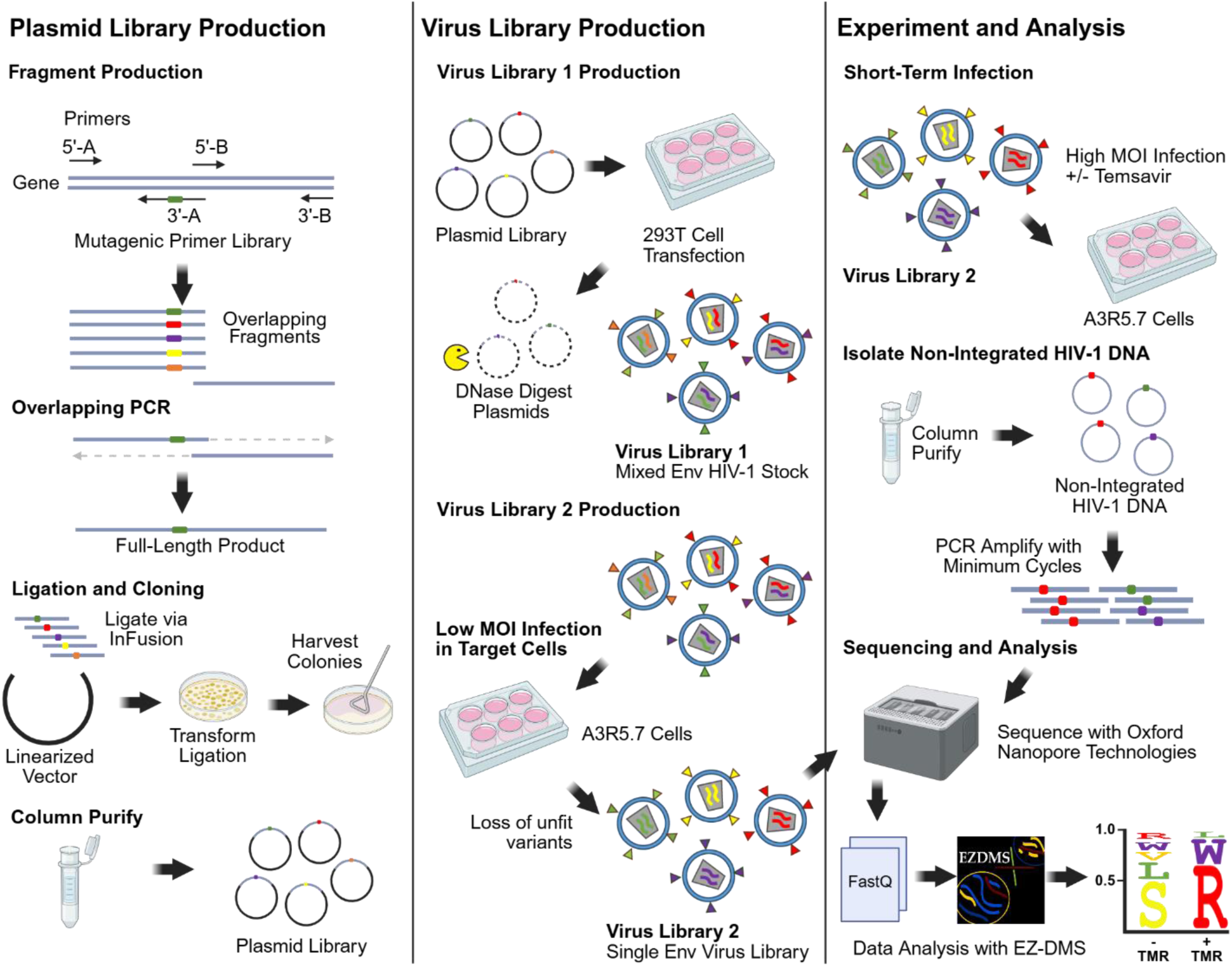
Outline of the EZ-DMS protocol

**(Step 1.1) Library construction** begins with the design of mutagenic primers with degenerate codons at the target site. To this end, we introduce the trinucleotide sequence NNK, where N is any nucleotide, and K is guanine (G) or thymine (T). This sequence encodes for 32 different codons, including all amino acids and a single (amber) stop codon. The amplicon region is first identified, and two flanking primers are designed for In-Fusion Cloning into the vector, each containing exactly 15 bp overlaps with the vector.

**(Step 1.2) Fragment Production:** Two fragments are produced, which will be ligated by overlap extension PCR. Fragment A is generated from the 5’-A flanking primer and a 3’-A primer (60 bp) containing the NNK site. The 3’-A primer also contains a 30 bp overlap with Fragment B. Fragment B is generated with a 3’-B primer that flanks the gene, and a 5’-B primer with the 30 bp overlap with Fragment A. For incorporation of two NNK sites, 3 fragments may need to be produced depending on the distance between the sites. To allow multiplexing of samples in the sequencing reaction, unique barcodes are added to any of the primers. For our application, we included 2 synonymous mutations in a 10 bp region near the NNK site of primer 3’-A.

**(Step 1.3) Overlap extension PCR** is employed to generate a full-length product, followed by several rounds of PCR with the flanking primers and subsequent gel purification. All PCR reactions are optimized to reduce amplification bias through increased template and primer concentrations, a reduced number of cycles, and are performed in triplicate with pooling of the products.

**(Step 1.4) Ligation and Cloning**: The vector is linearized via PCR to delete Env using a primer that contains the AfeI restriction site and a primer that contains the SmaI site. The two primers contain an overhang that is used for recircularizing the Env-deleted amplicon (for propagation in bacteria and storage) via the In-Fusion cloning method. To generate the vector for Env cloning, the Env-deleted plasmid is digested by the above restriction enzymes. To clone the Env amplicon library into the linearized vector, we also use the In-Fusion method, and the product is transformed into the chemically competent Zymo Mix&Go! DH5α cells. Transformed cells are plated directly onto agar without heat shock, outgrowth, or recovery. Combining In-Fusion cloning with no-outgrowth, chemically competent cells significantly increased the yield of correctly ligated products (**Figure 2**). It is important that the transformation produces sufficient colonies so that each amino acid at the site of interest is sampled. Each unique NNK combination has a 1/32 chance of occurring per colony. With independent sampling, the 95% joint probability for selecting enough colonies to have every amino acid present, accounting for synonymous NNK codons, is 172 colonies for single NNK sites and 8,128 colonies for two NNK sites (43). To minimize bias in library distribution that may occur during replication in broth media, plasmid libraries are recovered from the plates by scraping the colonies into liquid medium, followed by a short recovery period of approximately 2 hours. Plasmid libraries are then purified using either mini- or midi-prep kits.

**Figure 2:**
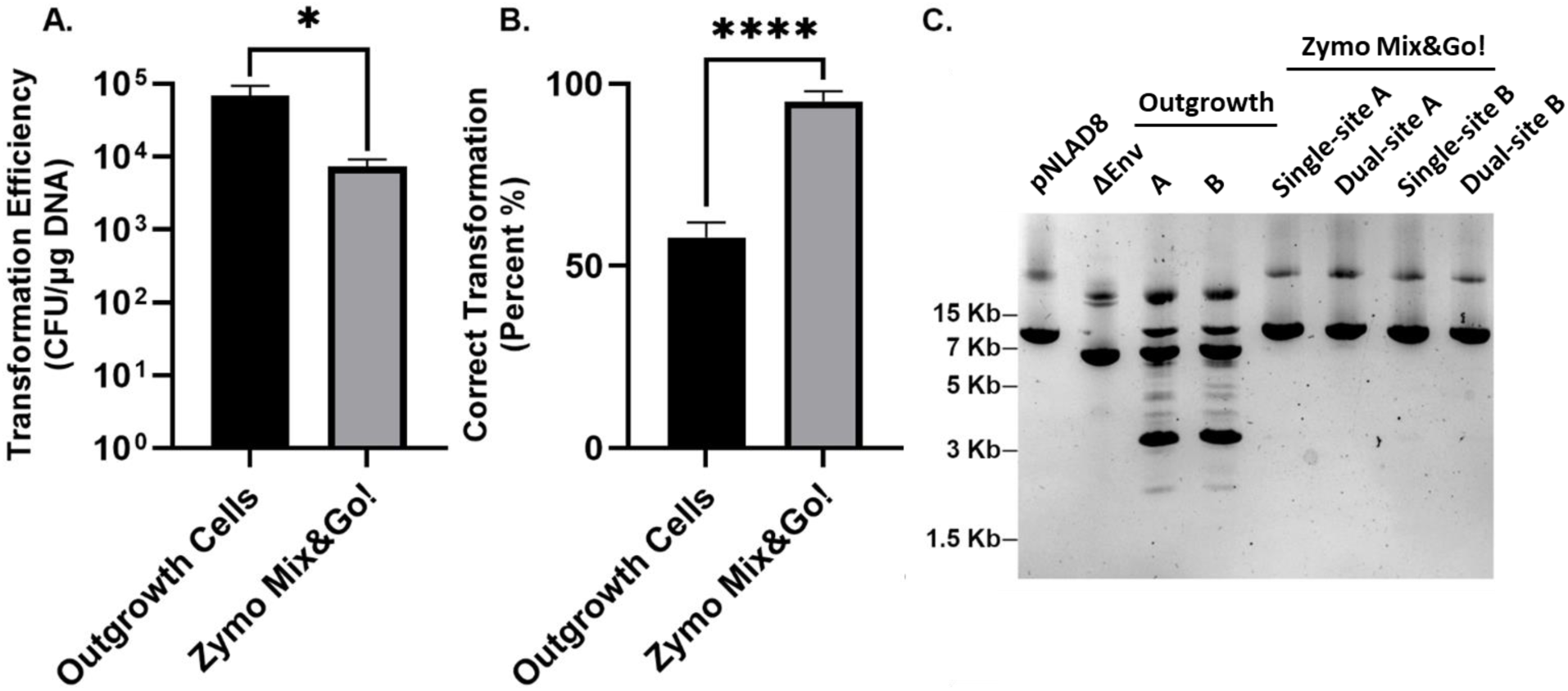
Transformation efficiency and accuracy in Stellar chemically competent (Outgrowth) cells and Zymo Mix&Go! cells. (**A**) Transformation efficiency using the two methods. (**B**) Percent colonies that contain the correct insert size, as determined by colony PCR screening. Bars represent the mean and standard error of the mean (SEM) from nine independent transformations. Statistical significance of the differences between conditions was calculated using an unpaired t-test: *, P < 0.01; ****, P < 0.0001. (**C**) The gel shows from left to right: **(i)** Full-length pNLAD8 plasmid, **(ii)** Env-deleted vector for ligation, **(iii)** Two single-site plasmid libraries harvested from outgrowth cells, **(iv)** Two replicates of libraries (A and B) from single-site and dual-site DMS. All libraries were independently constructed from the same cloning vector.

### (2) Virus Library Production

**Virus Library 1** is generated by transfecting the plasmid library into HEK 293T cells using JetPrime reagent (PolyPlus Inc.) or another suitable transfection protocol. Cells usually uptake more than one plasmid when transfected (44, 45). Thus, due to the nature of HIV-1, which contains two genomes per virion, the virions produced from this step predominantly contain two distinct genotypes. To obtain a virus library with a single genotype per virion, an additional passage of the virus is performed at a low multiplicity of infection (MOI) of 0.003 (for single NNK-site libraries) or 0.006 (for dual-site DMS libraries). For our application, the target cells used for infections were the T-cell acute lymphoblastic leukemia A3R5.7 cells (46, 47). These cells naturally express the HIV-1 receptor CD4 and coreceptor CXCR4, as well as the coreceptor CCR5 under selection of geneticin. The resulting **Virus Library 2** is harvested. If sufficient diversity is not achieved for the virus stock, this may be increase by ultracentrifugation over a sucrose cushion (48).

### (3) Experiment and Analysis

In this study, we tested our libraries by performing a **Short-Term Infection** in A3R5.7 cells (**Figure 1**) in the presence of the entry inhibitor temsavir (49–51). This inhibitor interacts with Ser at position 375 and Met at position 426 of Env, according to standard HXBc2 numbering (52). To test the effects of mutations at these sites on resistance to temsavir, we performed DMS as described above for positions 375 and 426, as well as dual-site DMS for both positions. Each Virus Library 2 was used to infect the cells for 16 hours, after which the cells were harvested, and the reverse transcribed (non-integrated) viral DNA was isolated from them using QIAprep Spin Miniprep kit (Qiagen). Three sequencing reactions of the region containing the NNK sites were submitted for each experiment: **(i)** The plasmid library, **(ii)** Virus Library 2, after extraction of viral RNA and reverse transcription to produce cDNA, and **(iii)** The cell-derived pre-integrated viral DNA collected following the short-term infection. For **Sequencing and Analysis,** PCR products were column-purified and sequenced via Oxford Nanopore technology. As a sequencing service provider, we used PlasmidSaurus, which provides up to 12,000 raw reads per sample, allowing samples from different experiments to be multiplexed.

The generated FASTQ files are then directly uploaded to the graphical user interface-based software EZ-DMS, which determines for each DMS site the amino acid frequency and calculates the. The following calculations to determine preferences, which are based on those previously described by the Bloom Group (12, 32), are automatically performed by the EZ-DMS software suite, which can be found at: https://github.com/haimlab/EZDMS, To calculate the enrichment ratio (𝜙) for each amino acid 𝑎 at position 𝑝, we used:

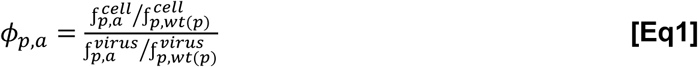

where 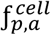 and 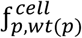 are the frequencies in the cell lysate of amino acid 𝑎 or the wild-type amino acid for position 𝑝, 𝑤𝑡(𝑝), and 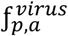 and 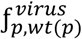 are the frequencies of these forms in the input virus sample used for infection. The amino acid preference (𝜋) for each amino acid is then calculated as:

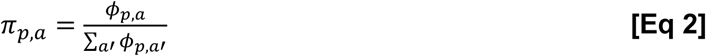

where the ∑_𝑎′_ 𝜙_𝑝,𝑎′_ is the sum of enrichments for all amino acids 𝑎′ at position 𝑝.

## EXPECTED RESULTS

Herein, we present the results of two independently constructed libraries for HIV-1 Env, generated from separately ordered primers and performed in triplicate. Both experiments utilized a single-site library at Env position 375 and a dual-site library at positions 375 and 426. For single-and dual-site experiments, the plasmid libraries were compared to a hypothetical distribution (**Figure 3A** and **3D**). Additionally, independent replicates of the plasmid libraries and virus libraries (replicates A and B) were compared (**Figure 3B, 3C, 3E** and **3F**).

**Figure 3:**
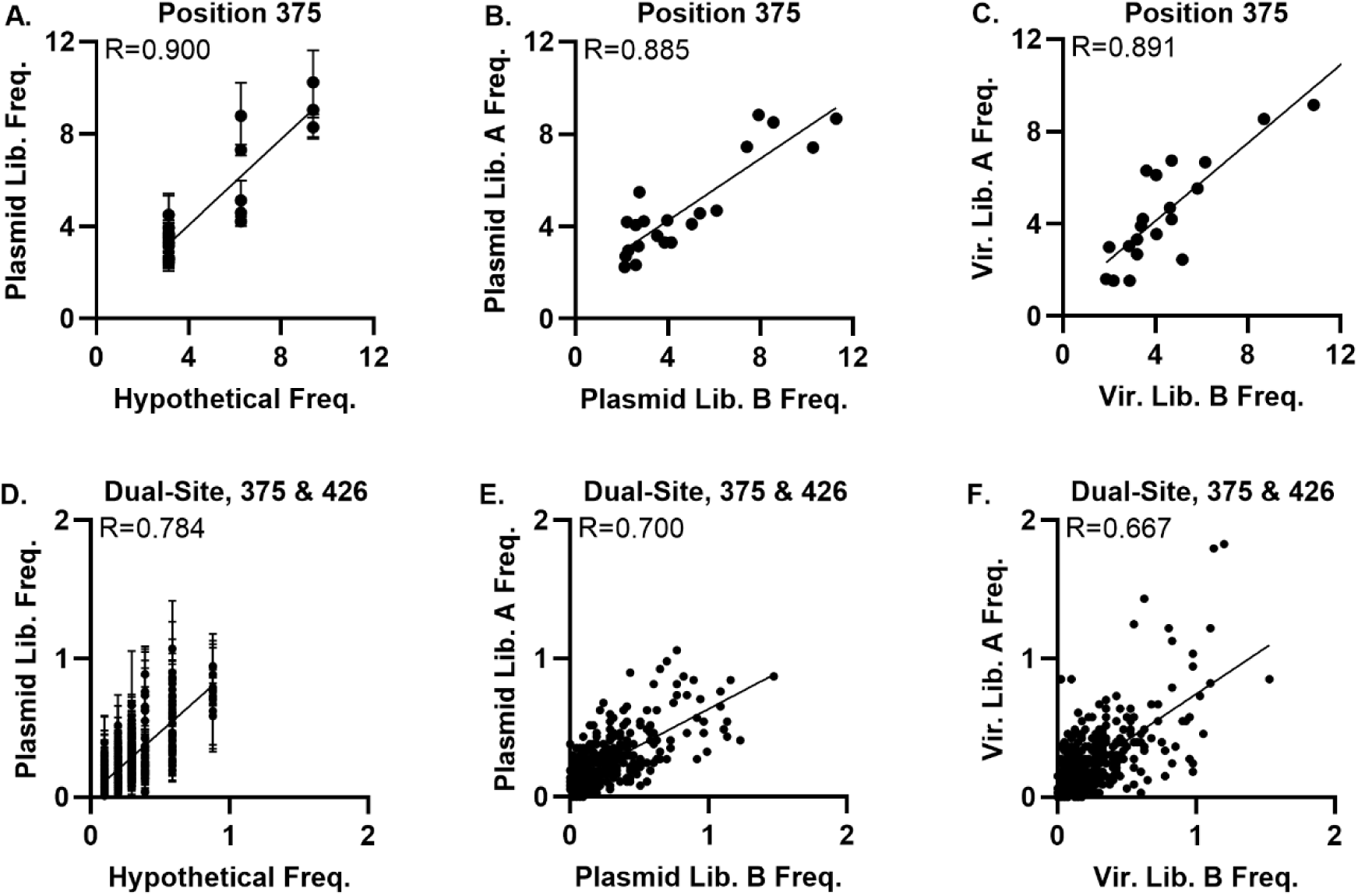
Plasmid and virus library validation. The data show correlations for the single-site (**A-C**) and dual-site (**D-F**) DMS libraries. Frequencies of amino acids in the two replicates of the plasmid libraries are compared with their expected frequencies (hypothetical) (**A, D**). Error bars represent the SEM among the two replicates of the libraries. The correlation between independent replicates of the plasmid libraries is shown in panels **B** and **E**, and between the Virus Library 2 replicates in panels **C** and **F**. The Pearson correlation coefficient (R) is shown for each comparison.

Next, we evaluated the effect of temsavir at the IC_99_ (250 nM) of this agent for HIV-1 AD8 Env on inhibition of entry by virus containing DMS libraries at positions 375 and 426. The A3R5.7 cells were used as targets for the short-term infection (**Figure 4**). In the absence of inhibitor, several amino acids exhibited low fitness (**Figure 4A** and **4C**). Excellent reproducibility was observed between the two independent replications (see error bars in **Figure 4A** and **4C** and correlations in **Figure 4E**). In the presence of 250 nM temsavir, selection at both positions was observed against the sensitive wild-type amino acids and in favor of known resistant mutants such as Met, His and Trp at position 375, and Leu at position 426 (**Figure 4B** and **4D**) (53). Novel resistant substitutions (Phe, Leu and Tyr at position 375) were also observed (**Figure 4B**). Interestingly, position 426 only exhibited a dramatically higher preference for Leu in the presence of temsavir relative to all other amino acids (**Figure 4D**). Indeed, the M426L mutation is the most commonly observed *in vitro* and in clinical samples from treated individuals (53, 54).

**Figure 4:**
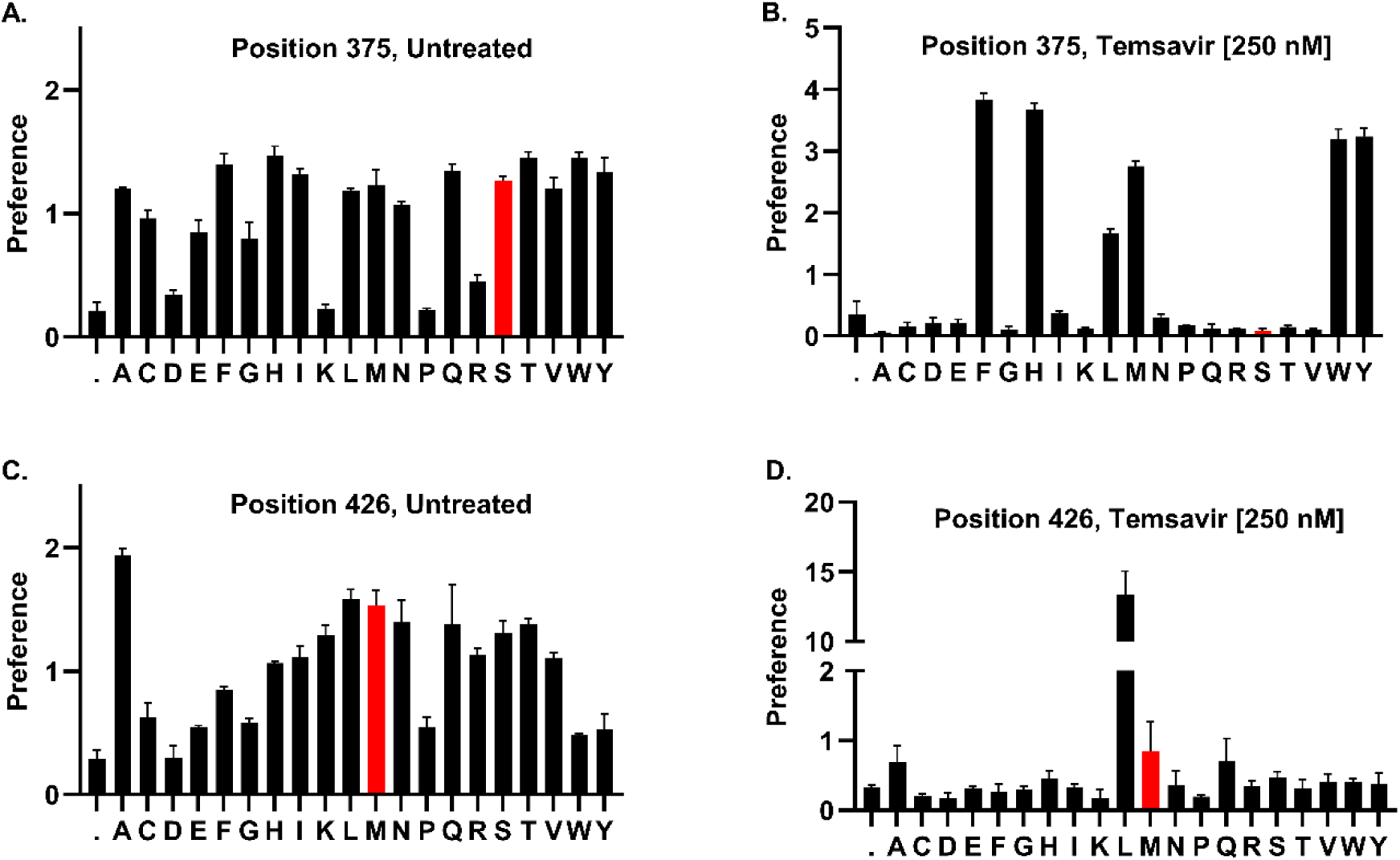
Amino acid preferences for single-site DMS at positions 375 and 426 in the absence and presence of temsavir. (**A,C**) Amino acid preferences at Env positions 375 or 426 in untreated cells. (**B,D**) Amino acid preferences at positions 375 and 426 in cells treated with 250 nM temsavir. The wild-type amino acid at each position is colored in red.

We also evaluated the fitness of dual-site DMS for the above two positions after a single round of infection of A3R5.7 cells in the absence and presence of 250 nM temsavir. Co-occurrence of mutations at the two sites have been observed in clinical trials (54). A total of 441 possible combinations of amino acids (and a stop codon) at Env positions 375 and 426 exist, of which 383 were observed in the input virus library. All combinations detected in the presence and absence of temsavir are shown in **Figure 5A**. We compared the frequencies of amino acids at position 375 in these data with their frequencies in the single-site DMS results and observed strong correlations (**Figure 5E** and **5F**). For the library with a single DMS site at position 426, a single amino acid (Leu) was dominant in the temsavir-treated samples (**Figure 4D)**. However, in the dual-site library, a greater diversity of amino acids at this position was observed in the presence of temsavir, potentially due to epistatic effects with variants at position 375 (**Figure 5C**).

**Figure 5:**
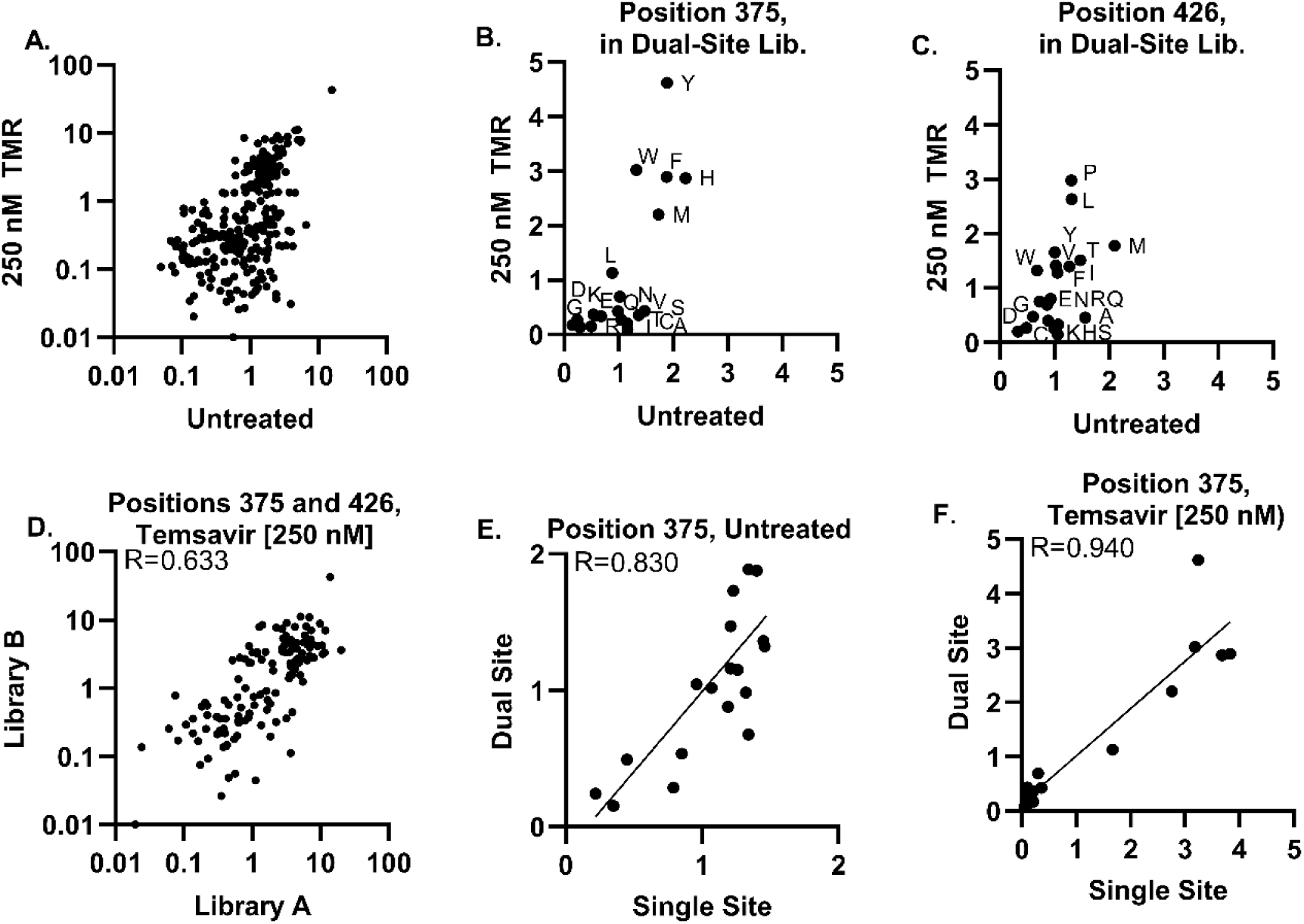
Amino acid preferences after a single round of infection using a dual-site DMS library at Env positions 375 and 426. Experiments were performed in the absence and presence of 250 nM temsavir (TMR). (**A**) Relationship between preferences for all amino acid combinations detected in the presence and absence of temsavir. **(B,C)** Relationships between the sum of all preference values for the 20 amino acids at position 375 or 426 in the dual-site DMS library. (**D**) Correlation between frequencies of amino acid combinations at the two sites in two replicate experiments. **(E, F)** Relationship between the frequency of the 20 amino acids at position 375 in the single-site DMS and the sum of all combinations in the dual-site DMS that contain them. The Pearson correlation coefficient is shown for each comparison.

## DISCUSSION

The EZ-DMS protocol and software suite can be used to address questions related to the evolutionary landscape of proteins from diverse virus types. Given the setup of the protocol, the requirements for viruses to be tested are: (**1**) The virus is replication competent. (**2**) Transformation efficiency is sufficient to produce enough colonies for the required library diversity. (**4**) The virus, either naturally or through low MOI passage, packages one genotype per virion. (**5**) The experimental system can separate the genetic material of infectious virus from non-infectious virus.

### Applications of the system include

#### 1. Mapping the fitness profile of viral proteins

In these experiments, infections are conducted in the absence of an inhibitor. Virus stocks are used to infect a population of target cells, most commonly by multiplexing libraries that span multiple positions of the target protein. The output is the relative infectivity of all amino acid variants of a protein compared with the wild-type form. When conducted using target cells that express alternate receptors for the virus, such tests can reveal the potential of the virus to alter cell tropism, and the specific changes required to achieve it (55, 56). Fitness experiments can also be performed to compare the profiles of specific positions in distinct virus isolates to gain a better understanding of mutation entrenchment. This phenomenon describes the changes in preference for amino acids at sites of interest due to accumulation of accessory mutations during divergence of the two strains (57).

#### 2. Determining the escape potential of the virus from therapeutics

Infections are conducted in the absence and presence of an inhibitor to measure the relative resistance and fitness of all variants. Such analyses have been conducted for Env-targeting antibodies such as PGT151 (58) and 1-18 (34). Here, we describe the profiling of escape mutations from the HIV-1 entry inhibitor temsavir (BMS-626529) (59). This FDA-approved agent binds to the CD4-binding of pocket of Env and prevents activation of the entry cascade by the CD4 receptor on cells (60). These experiments describe the genetic barriers to the development of resistance to the therapeutic.

#### 3. Identify epistatic interactions across proteins

Epistasis describes a relationship between two or more sites of a protein, whereby the amino acid occupancy at one site impacts the fitness profile (mutational tolerance) of another. Epistatic relationships within the different HIV-1 proteins can impact their escape patterns from inhibitors, as we have shown (6, 61), and are still poorly defined. To better understand such interactions, DMS can be applied. One approach involves introduction of a substitution at a site 𝑖 and analysis of the fitness profile at site 𝑗 for the 𝑖-mutant and wild-type forms. Differences between the observed profiles can capture the effect of the position 𝑖 occupancy on the fitness profile of position 𝑗. To gain a more comprehensive understanding of the effects, one can perform two-site combination tests, whereby degenerate codons are introduced at both positions 𝑖 and 𝑗. In this manuscript, as an example, we describe the introduction of degenerate codons at positions 375 and 426 of Env, which are the most commonly occurring resistance mutations to temsavir in treated subjects (50).

## ACKNOWLEDGEMENTS

This work was supported by National Institutes of Health (NIH) grant R01 AI170205 to HH. The funders played no role in study design, data analysis, *in vitro* data acquisition or analysis, or the decision to publish this work.

